# Cap-Independent Translation Regulation of VEGF Using CRISPR-dCas13d

**DOI:** 10.64898/2026.09.04.749420

**Authors:** Mohammad Lutful Kabir, Sineth G. Kodikara, Najmah Al Ramel, Janan Alfehaid, Soumitra Basu, Hamza Balci

## Abstract

Vascular endothelial growth factor (VEGF) is a central driver of pathological angiogenesis in cancer and retinal disease. A G-quadruplex (GQ) structure that forms in an Internal Ribosome Entry Site (IRES) of VEGF was proposed to recruit the 40S ribosome and initiate cap-independent translation under hypoxia. We demonstrate that blocking this GQ with dCas13d represses VEGF translation by up to 5-fold, without impacting mRNA abundance, under hypoxia. Similar levels of repression were attained in HeLa and human umbilical vein endothelial cells (HUVECs) using immunofluorescence and western blot assays, as well as a bicistronic dual-luciferase experiment. Single molecule Förster resonance energy transfer (smFRET) and fluorescence enhancement assays demonstrated destabilization of the GQ by CRISPR-dCas13d. We also tested the functional consequences of this regulation using tube formation assay in bovine aortic endothelial cells (BAOECs), which demonstrated consistent results with VEGF expression levels. Our study demonstrates the feasibility of using CRISPR-dCas13d as a sequence-specific and transient translation regulator for a large group of genes that contain translationally-active secondary structures.

**Graphical Abstract:** Graphical Abstract
Target the VEGF-GQ located within IRES-A with CRISPR-dCas13d modulates VEGF translation.

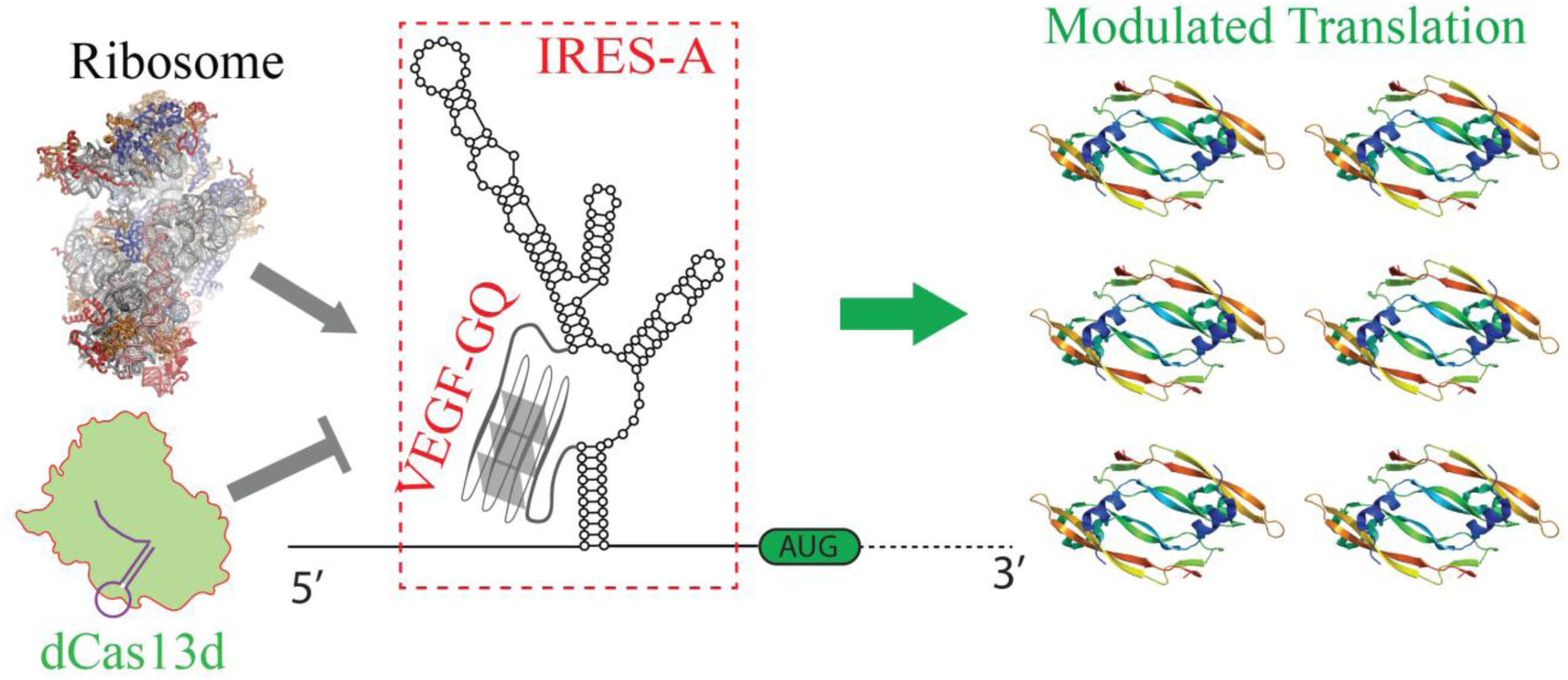

## Introduction

Vascular Endothelial Growth Factor (VEGF) is a homodimeric glycoprotein that plays crucial roles in angiogenesis and promotes the proliferation and migration of endothelial cells, which line blood vessels ^1,2^. VEGF stimulates the formation of new capillaries, helping to supply oxygen and nutrients to tissues ^1,3,4^. When oxygen levels are low, VEGF production, in particular its VEGF-A isoform, increases to enhance blood supply and adapt to hypoxia ^5,6^. VEGF is essential for normal physiological processes, and it is overexpressed during embryonic development and wound repair; however, its overexpression is also linked to various diseases, including cancer facilitating tumor growth and metastasis and is a master regulator of tumor angiogenesis and vascular disorders ^7^. Excessive VEGF can lead to abnormal blood vessel growth in the eye, causing vision loss, such as age-related macular degeneration and diabetic retinopathy ^8^. Repressing VEGF can help ameliorate these conditions by starving tumors of the required oxygen ^9–11^.

Targeting VEGF is a cornerstone strategy in angiogenesis research and therapy; therefore, VEGF has been targeted at different stages with various approaches including (i) monoclonal antibodies (e.g., bevacizumab) and decoy receptors (e.g., aflibercept) that prevent VEGF from activating its receptors ^12^; (ii) small-molecule tyrosine kinase inhibitors (e.g., PDGFR, FGFR) that block downstream VEGF receptor signaling^13^; (iii) transcriptional regulation of VEGF by inhibiting activating transcription factors (e.g., HIF-1α) ^14^; (iv) post-transcriptional regulation of VEGF mRNA by RNA interference ^15^ or anti-sense oligonucleotides ^16^. In this study, we utilized clustered regularly interspaced short palindromic repeats (CRISPR) and associated (Cas) proteins to repress human VEGF (hVEGF) expression by targeting a G-quadruplex forming sequence (QS) located in its 5′ untranslated region (5′-UTR). For brevity, henceforth we will refer to this sequence as VEGF-rQS.

CRISPR functions as an RNA-driven adaptive immune system that protects bacteria and archaea against invading bacteriophages ^17,18^; however, nuclease-deficient Cas (dCas) proteins have been adapted to regulate gene expression. For example, dCas9 has been adapted to target genomic DNA ^19,20^ to regulate transcription in a sequence-specific manner. Similar mutations have been introduced in Cas13d, an RNA-targeting Cas protein, to render it nuclease-inactive. Here, we used dCas13d from the bacterium *Ruminococcus flavefaciens* ^21,22^, where R239A, H244A, R858A, H863A mutations are introduced into the two higher eukaryotes and prokaryotes nucleotide binding (HEPN) domains of Cas13d. This system has been used in various applications, including translation activation and repression ^23–26^. Using CRISPR-dCas13d, we targeted VEGF-rQS in order to regulate VEGF expression at the translation level.

The conventional process of translation initiation relies on the 5’-cap to coordinate the assembly of translation initiation machinery ^27^. In an alternative process, ribosomes are recruited to highly structured regions in the 5’-UTR, known as Internal Ribosome Entry Sites (IRES), which are particularly significant for viruses (including PV, EMCV, HCV, FMDV, Swine Fever virus, and HIV) ^28,29^. IRESs have also been discovered in eukaryotic systems, including in several growth factors with longer than normal 5′-UTR. The 5′-UTR of *hVEGF* includes IRES-A and IRES-B which are capable of independently initiating cap-independent translation ^30^. Under hypoxic conditions, VEGF expression is under the control of IRES-A which initiates translation of a secreted form of VEGF-A ^31^. IRES-A is 293 nucleotide (nt) long (745-1038 nt from 5’-end) and is located immediately upstream of the canonical translation start site (AUG at 1039 nt) (Figure 1). The ribosome typically scans along the mRNA for approximately 200 nt before locating the start codon. The VEGF-rQS is a 17 nt long segment (GGAGGAGGGGGAGGAGG, 774-790 nt) within IRES-A and promotes translation by facilitating ribosome recruitment under hypoxia ^32,33^. We used dCas13d to destabilize this GQ and inhibit ribosome recruitment to IRES-A in order to inhibit translation initiation.

**Figure 1.**
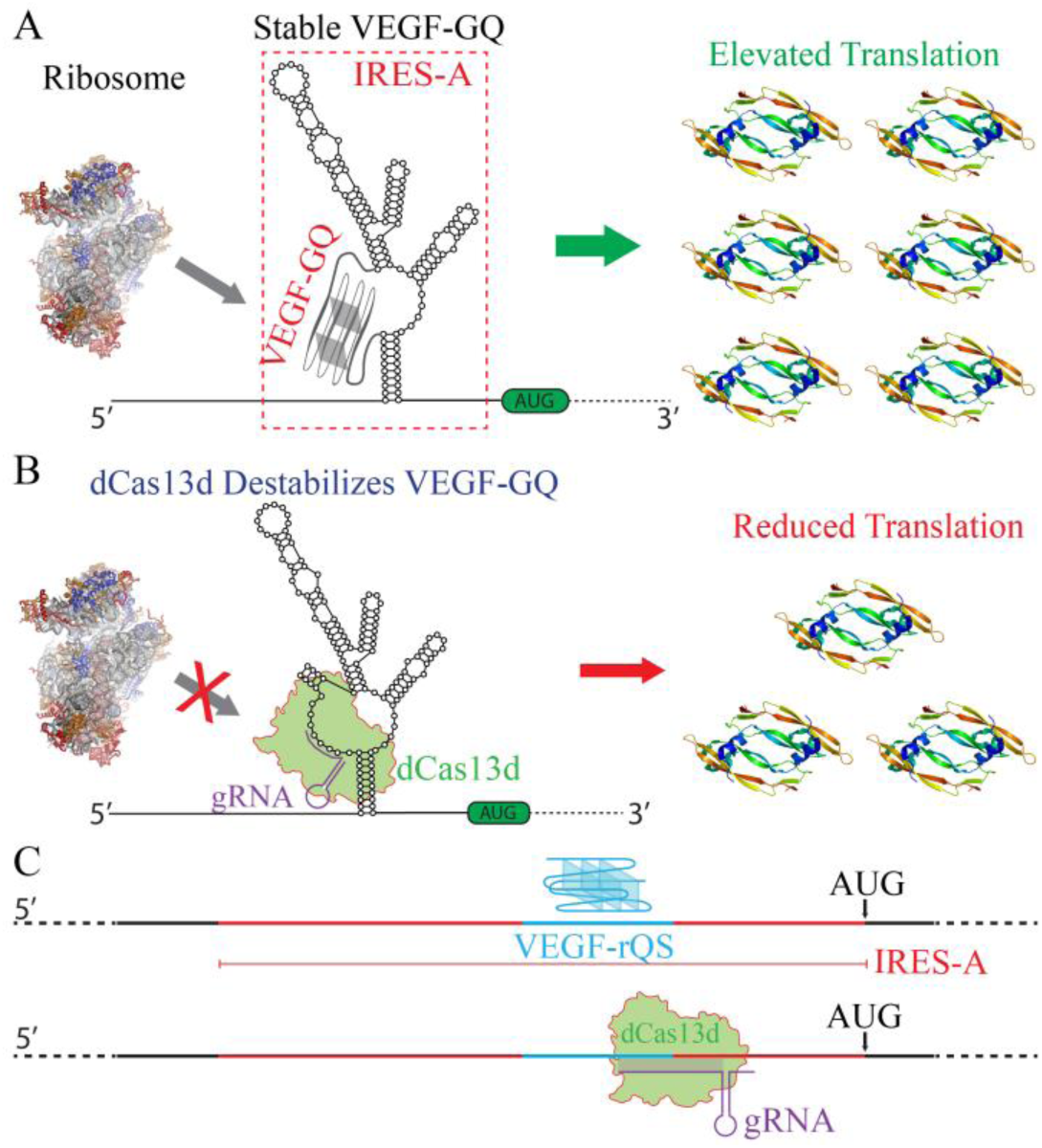
Schematics showing VEGF-rQS and IRES-A within the 5’-UTR. A) Cap-independent translation of VEGF-A via recruitment of the 40S ribosome to a GQ located within IRES-A. AUG is the translation start site. B) A schematic showing the hypothesis driving this study: destabilizing the GQ within IRES-A prevents ribosome recruitment and represses VEGF-A translation. C) Linearized version of 5’-UTR of VEGF, location of IRES-A, and CRISPR-dCas13d target site immediately upstream of the start codon.

The dCas13d based approach presented in this study has similarities but also significant differences from antisense oligonucleotides (ASOs), a prominent and mature strategy for translation regulation. ASOs are 12-30 nt long oligonucleotides and are often chemically modified for various purposes. They bind complementary RNA sequences and either promote double stranded RNA degradation via RNaseH-mediated cleavage or block ribosome binding when targeted to the translation start site ^34,35^. Degradation of mRNA has typically been the preferred mechanism for therapeutic applications as greater levels of repression in protein expression levels can be achieved ^36^. However, this raises concerns about the consequences of off-target effects as mRNA stability is impacted. In contrast, dCas13d does not promote RNA degradation and requires both guide RNA complementarity and binding of the dCas13d protein itself. This dual requirement may repress weak or transient interactions, thereby reducing off-target effects.

Genome-wide computational studies and high-throughput sequencing have identified several hundred thousand intramolecular putative GQ forming sequence (PQS) in the human genome ^37^. Telomeres and promoters, especially the immediate vicinity of transcription start site (TSS), are rich in PQS ^38–40^. About 50% of human genes contain a PQS within 1,000 nts upstream of TSS ^41^. PQS are more prevalent in promoters and UTRs of oncogenes and regulatory genes, such as transcription factors, compared to housekeeping genes ^42^ and are involved in transcription and translation regulation (17–21). Therefore, targeting the PQS with dCas13d promises to be a widely applicable and sequence specific method to regulate translation, unlike alternative approaches, such as small molecules or site directed mutagenesis, which lack sequence specificity or transience. In recent studies, we demonstrated the applicability of this approach at the transcription level by targeting a PQS in *tyrosine hydroxylase* (*TH*) and *c-Myc* promoters with CRISPR-dCas9 ^43,44^. In particular, we demonstrated that c-Myc mRNA and protein expressions can be repressed by up to 3-fold by targeting one site and up to 10-fold by simultaneously targeting two specific sites withing the vicinity of PQS with CRISPR-dCas9 ^44^. However, to our knowledge, such an approach, which relies on modulating GQ stability, has not been demonstrated at the translation level yet. This requires an RNA specific CRISPR system and a PQS that is involved in translation regulation ^45–47^. Cas13d is one of the most prominent RNA-targeting CRISPR systems. However, until the nuclease-dead dCas13d was engineered several years ago, gene regulation via Cas13 (formerly C2c2) relied on cleaving target RNA ^48,49^, which risked unwanted RNA degradation. Recent studies have demonstrated the feasibility of translation regulation using dCas13d or a fusion of dCas13d and a partner (such as elF4G) ^50–52^. While modulation of secondary structure forms the core of our approach, translation regulation in other studies has predominantly relied on blocking ribosome scanning or binding.

Despite supporting evidence from multiple studies about the role of VEGF-rQS in cap-independent translation ^32,33,53,54^, this sequence was proposed to be dispensable for cap-independent translation and stabilizing it with mutations and small molecules repressed translation ^45^. Therefore, it is important to establish the role of VEGF-rQS in translation without introducing sequence mutations or using small molecules, which is the goal of the current study.

## Materials and Methods

### Plasmid Constructions

The bicistronic plasmid (hVEGFbicis) containing Renilla and Firefly Luciferase genes and wild-type IRES-A sequence, including the G-quadruplex forming VEGF-rQS, was constructed as previously described ^32^. Renilla Luciferase is under the control of CMV promoter while Firefly Luciferase is under IRES-A control.

### RNA Preparations

All RNA oligonucleotide sequences are reported in Supplementary Table S1. The VEGF-rQS is a 17-nt-long segment (GGAGGAGGGGGAGGAGG, 774-790 nt) within IRES-A. The guide RNA (gRNA) strand and dCas13d protein were mixed together to form the ribonucleoprotein (RNP) complex. All guide RNA oligonucleotides were purchased from either Integrated DNA Technologies (IDT) or Dharmacon, and the dCas13d proteins (from *Ruminococcus flavefaciens* (casRX)) were purchased from MCLab. Guide RNA strands were purified via denaturing polyacrylamide gel electrophoresis (PAGE) with different percentages of gel solutions. Full-length products were visualized by UV shadowing, marked, and then excised from the gel. The RNA was harvested via the crush and soak method by tumbling the gel slice overnight at 4 °C in a solution of 300 mM NaCl, 10 mM Tris–HCl, and 0.1 mM EDTA (pH 7.4). Salt was removed by ethanol precipitation of the oligonucleotides twice, with two cold 70% (v/v) ethanol washes in between each precipitation. The oligonucleotides were dissolved in nuclease-free water and stored at −20 °C.

### Single Molecule FRET Experiments

A home-built prism type total internal reflection fluorescence microscope was used for single molecule experiments. The slides and coverslips were treated with 1M KOH followed by piranha etching. The slide surfaces were functionalized with amino silane solution followed by surface passivation with polyethylene glycol (PEG) over night. Additionally, the surfaces were treated with a second round of 333 kDa PEG to increase the density of the PEG brush and enhance the surface quality. The microfluidic chamber was created between a slide and cover slip with double-sided tape separating them. The chamber was washed with T50 buffer (10 mM Tris, 2 mM MgCl2 and 50 mM KCl) before adding streptavidin followed by addition of the biotinylated RNA-DNA hybrid constructs or RNA-DNA constructs mixed with the RNPs (preparation described below). The imaging was done with a custom-built C++ program ^56^ using an Andor Ixon EMCCD camera. We recorded short (15 frames) and long movies (1500 frames) for each studied condition and an image acquisition rate of 100 ms per frame. The data was analyzed in MATLAB, plotted and fitted in Origin.

Single molecule FRET experiments were performed on surface-immobilized partial duplex RNA-DNA hybrid constructs that have long RNA and a short complementary DNA. PAGE-purified 60-nt-long RNA strand (with a 5’-Cy3 label) and a 12-nt-long complementary DNA strand (with a 5’-biotin and 3’-Cy5) were hybridized. The RNA and DNA strands were annealed at a 3:1 molar ratio at 95 °C for 5 min in the presence of 150 mM KCl and 2 mM MgCl2, followed by slow cooling to room temperature. For reconstitution of ribonuclear protein (RNP) complexes, dCas13d was incubated with a 2-fold excess of gRNA at 37 °C for 30 min in RNP buffer (50 mM Tris-HCl, pH 7.5, 100 mM NaCl, 6 mM MgCl2, 1 mM DTT) ^57^. Then, the RNP complex was mixed with the annealed RNA-DNA hybrid construct with 1x working buffer (200 mM HEPES, 50 mM MgCl2, 1 M NaCl, 1 mM EDTA, pH 6.5). These complexes were diluted to 10-50 pM (in 150 mM KCl and 2 mM MgCl2), introduced to the microfluidic sample chamber, and incubated for 2-5 min, to achieve a surface density of ∼300 molecules per imaging area (∼50 µm x 100 µm). Excess or unbound molecules were removed from the chamber by washing the chamber with an imaging buffer (50 mM Tris-HCl (pH 7.5), 2 mM Trolox, 0.8 mg/mL glucose, 1% gloxy (0.1 mg/mL glucose oxidase and 0.02 mg/ml catalase), 0.1 mg/mL bovine serum albumin (BSA), 2 mM MgCl2, and 150 mM KCl).

### Fluorescence Enhancement Assay

N-Methylmesoporphyrin IX (NMM) was purchased from Thermo Fisher Scientific and diluted in 1 mM DMSO. Any remnant secondary structure in RNA oligonucleotides (VEGF-GQ or guide RNA strand) was reset by heating them to 95 °C in the presence of a buffer containing 150 mM KCl and 2 mM MgCl2 [pH 7.5]. A gradual cooling to room temperature allows VEGF-GQ to fold. The dCas13d-gRNA RNP complexes (100 nM, RNP1-4) were incubated with folded VEGF-GQ RNA strand (50 nM) at 37 °C for 30 min. Then, NMM (150 nM) and VEGF-GQ or VEGF-GQ + RNP complexes (100 µl) were mixed in a 96-well plate with black walls and transparent bottom. Control wells that include only NMM or NMM mixed with non-G-quadruplex RNA were also prepared. The plate was incubated in the dark for 15–30 min to allow binding interactions to occur. Fluorescence emission spectra (500-700 nm) were recorded using an Agilent BioTek Synergy Neo2 plate reader with an excitation wavelength of 399 nm. The data were analyzed by comparing the fluorescence enhancement of the NMM emission under different conditions.

### Immunofluorescence Assay

Immunofluorescence assay was used to determine VEGF protein expression ^59^. HUVECs were maintained in endothelial cell growth medium (EGM-2) with high glucose supplemented with 20% fetal bovine serum (FBS) and 1% antibiotics (streptomycin and penicillin) at 37 °C in 5% CO2 in a humidified incubator. HUVECs were treated with Sodium Nitroprusside (SNP at 100 µm) for 24 h to induce hypoxic conditions ^60,61^ and with different RNP (30 nM) complexes to investigate their impact on VEGF translation under hypoxia. Following fixation in paraformaldehyde, the treated and untreated cells were rinsed in phosphate buffer saline (PBS) buffer containing Tween-20 (0.05 %), prior to Triton X-100-mediated permeabilization. They were then blocked for 1 h with bovine serum albumin (1 %), rinsed three times in the rinsing solution, and incubated with anti-VEGF-A primary antibody (diluted 1:100) for 24 h, followed by rinsing and incubation for 1 h with secondary antibody labeled with Alexa Fluor 488. This was followed by rinsing and staining with 4’, 6-diamidino-2-phenylindole dihydrochloride (DAPI). After rinsing, the cells were imaged with scanning confocal microscopy, and images were analyzed and quantified with ImageJ.

### Luciferase Assay

HeLa cells were grown in 96-well plates in Dulbecco’s Modified Eagles Medium (DMEM) supplemented with 10% fetal bovine serum and the antibiotics streptomycin and penicillin at 37 °C in 5% CO2 in a humidified incubator. Approximately 70% confluent HeLa cells were transfected with vectors (hVEGFbicis) using Lipofectamine 2000 (Invitrogen) according to the manufacturer’s protocol, and those wells were treated with different RNPs (30 nM) complexes. 24 h after transfection and treatment, Renilla (RL) and Firefly (FL) luciferase activities were measured using a Dual-Glo Luciferase assay system (Promega) as per the manufacturer’s supplied protocol on an Agilent BioTek Synergy Neo2 plate reader.

### Quantitative RT-PCR Assay

HUVECs were maintained in endothelial cell growth medium (EGM-2) with high glucose supplemented with 20% fetal bovine serum (FBS) and 1% antibiotics (streptomycin and penicillin) at 37 °C in 5% CO2 in a humidified incubator. HUVECs were grown in 12-well plates at 37 °C in 5% CO2 in a humidified incubator for 24 h. These cells were treated with SNP and different CRISPR–dCas13d RNP complexes using TransIT-X2 transfection reagents (Mirus Bio, USA) when cells were ∼70% confluent. After the treatment, the cells were incubated for 24 h. The total RNA from transfected cells was extracted using the Trizol reagent following a previously optimized protocol. The complementary DNA (cDNA) was synthesized with 500 ng of total RNA using cDNA Super Mix (Quanta Biosystems, USA). Quantitative reverse-transcription polymerase chain reaction (RT-qPCR) assay was performed to quantify the endogenous VEGF-A messenger RNA (mRNA) level using specific primers and SYBR Green PCR Master Mix kit (Quanta Biosystems) on an Eppendorf Mastercycler RealPlex2 Sequence Detection System. The relative fold change in VEGF-A expression was determined using the Livak method ^62^.

### Western Blot

Total proteins were extracted from the HUVECs with RIPA buffer (Santa Cruz) 24 h after transfection with different CRISPR–dCas13d RNP complexes. 50 μg of protein lysate was separated by 15% SDS−PAGE, then transferred to the polyvinylidenfluoride (PVDF) membrane. Subsequently, they were blocked in 5% non-fat milk in PBS + 0.1% Tween-20. Then, the blotted membranes were incubated overnight at 4 °C with primary antibodies of anti-VEGF-A (1:500) (Rabbit mAb #50661, Cells Signaling Technology, USA). GAPDH was used as a loading control (G-9, sc-365062) antibody at a 1:1000 dilution. Horseradish peroxidase-conjugated goat anti-mouse IgG (sc-2005) was used as a secondary antibody at a 1:1000 dilution. Proteins were visualized by Western Blotting Luminol Reagent (sc2048) in ChemiDoc-ItTS2 Imager.

### Endothelial Tube Formation Assay (ETFA)

Bovine Aortic Endothelial cells (BAOEC), primary bovine endothelial cells, were purchased from CELL Applications, Inc (B304-05). These cells were cultured in Bovine Endothelial Growth Medium (B211-500) in a sterile biological safety cabinet.

In order to create networks in the ETFA, BAOEC 1x10^5^ cells/well on Matrigel (250 µl/well) were seeded into a 24-well plate and left for 24 hours at 37 °C with 5% CO2 (28). Cells within the well plate were treated with different CRISPR-dCas13d complexes to induce or suppress angiogenesis, while the control well was treated with the transfection reagent only. For hypoxia studies, cells were treated with Sodium Nitroprusside (SNP, 100 µM) to induce hypoxic conditions.

Photographs were taken at different time points using an Agilent BioTek Cytation 5 inverted microscope. Images were analyzed using ImageJ software using a program developed for ETFA (29). The "Angiogenesis Analyzer" for ImageJ (30) is extended by this plugin, developed in ImageJ’s macro language.

## Results and Discussion

Fig. 1 shows schematics of the hypothesis driving this study. The first step towards this goal is establishing folding of the GQ in VEGF-rQS and its destabilization by targeting its vicinity with CRISPR-dCas13d. To address this issue, we conducted single molecule *in vitro* assays, including mass photometry and single-molecule Förster resonance energy transfer (smFRET) experiments. The goal of the mass photometry experiments was to establish the feasibility of complex formation between dCas13d, gRNA and the relevant segment of the mRNA using our incubation protocol. The expected molecular weight of dCas13d is ∼112 kDa, while those of mRNA (60 nt) and gRNA (53 nt) constructs are in the 18-20 kDa range. Fig. 2A shows example data illustrating a peak at 104.6±0.6 kDa in case of dCas13d, a new peak at 125.8±2.0 kDa when dCas13d and gRNA are mixed together, and upward shift in mass to 140.6±2.0 kDa when dCas13d, gRNA, and mRNA are mixed together. The error bars are standard error of Gaussian function fits while the corresponding standard deviations are 19.4 kDa, 24.5 kDa, and 26.0 kDa, respectively. These results are within one standard deviation of expected masses of complexes.

**Figure 2.**
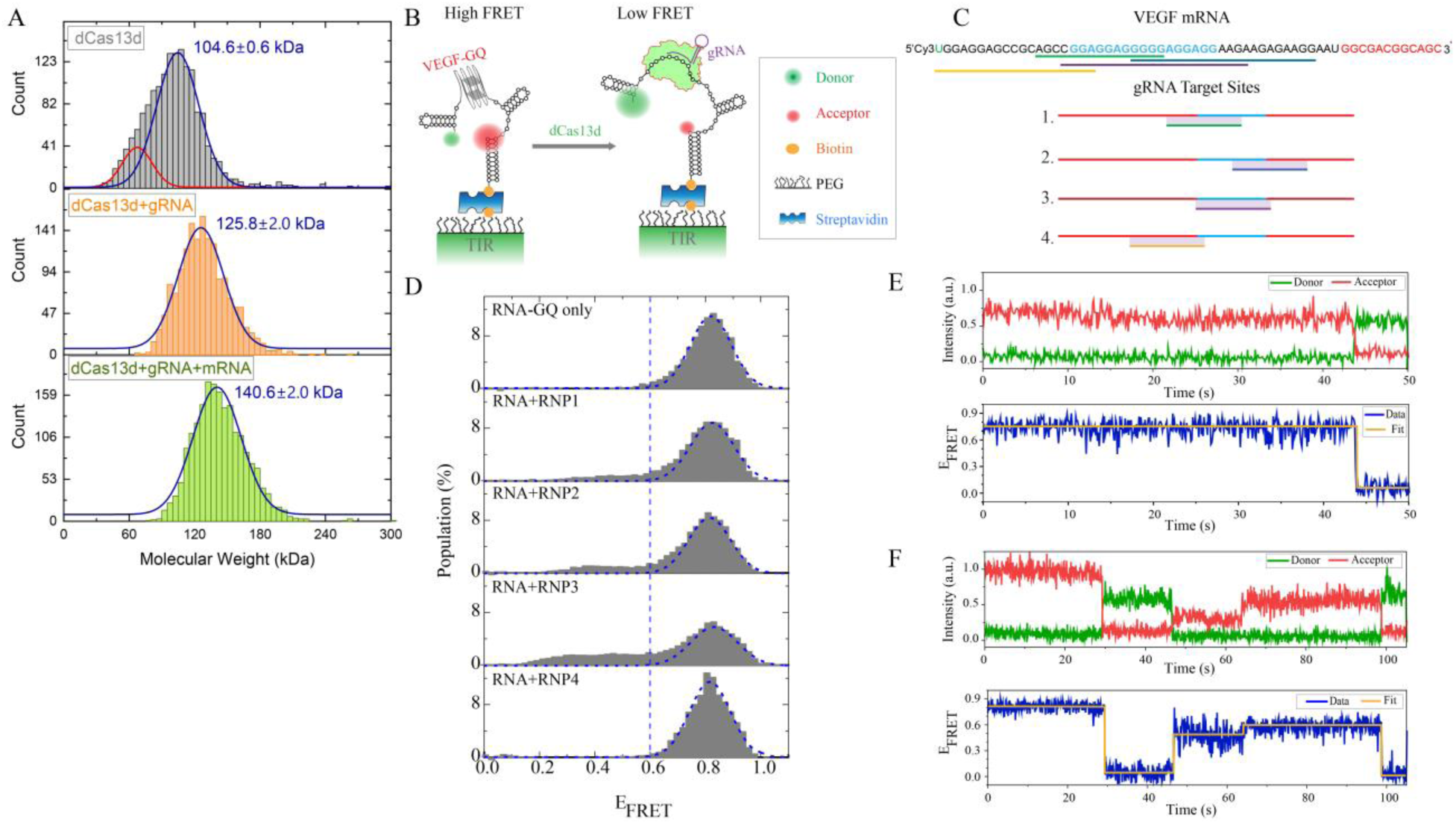
Single molecule mass photometry and FRET measurements. (A) Mass photometry illustrate complex formation between dCas13d, gRNA and mRNA under our incubation protocol. The relevant molecular weights are 110 kDa for dCas13d, 18 kDa for gRNA and 20 kDa for mRNA. The measured vs. expected molecular weights are 104.6±0.7 kDa vs. ∼112 kDa for dCas13d, 125.8±2.0 kDa vs. ∼130 kDa for dCas13d+gRNA, and 140.6±2.0 kDa vs. ∼150 kDa for dCas13d+gRNA+mRNA. The error bars are standard error of the Gaussian function fits. The small population in dCas13d case (top panel, red peak at 67.4±1.5 kDa) is due to a low mass background typically observed in mass photometry measurements or an impurity population. (B) Schematic of smFRET assay. The donor-acceptor fluorophores are placed such that the folded GQ results in high FRET while destabilization of the GQ with CRISPR-dCas13d complexes (RNPs) results in lower FRET. The illustrated hairpins are for demonstration purposes and do not represent the secondary structure of this truncated RNA construct. (C) Schematic of the four gRNA target sites, where the top panel shows the sequences targeted by these gRNA and the numbers 1-4 in the bottom panel refer to target sites for RNP1-4, respectively. (D) SmFRET histograms showing a single high FRET peak in the RNA construct before introducing the RNPs and development of a lower FRET population after introducing the RNP1-3. A total of 416 molecules were included in the histograms. (E-F) Example smFRET traces showing static (E), or dynamic (F) complex formation. The fraction of each type for different cases is given in the text. Additional sample traces are presented in Fig. S2.

In smFRET experiments, a 60 nt long mRNA, which includes the VEGF-rQS and neighboring sequences, was immobilized on the surface (Fig. 2B-C). We first identified the FRET level for folded GQ under stabilizing ionic conditions (150 mM KCl). These data (RNA-only) showed a single peak at FRET efficiency EFRET=0.80 (Fig. 2D, top panel). We then repeated these measurements by adding CRISPR– dCas13d ribonucleoprotein (RNP) complexes: RNP1, RNP2, RNP3 or RNP4 which target different sequences on the VEGF mRNA construct (Fig. 2C). The target sites for RNP1-3 overlap the VEGF-rQS; therefore, we expect the GQ to be destabilized and the distance between donor and acceptor fluorophores to significantly increase (Fig. 2B), which would result in a population at lower EFRET. RNP4 overlaps with only one of the five G-repeats in VEGF-rQS; therefore, it does not necessarily inhibit GQ formation. Control experiments in which gRNA were introduced without dCas13d did not show any change in the distributions (Fig. S1). Fig. 2D illustrates confirmation of these ideas where a population at lower EFRET and a reduction in the population of EFRET=0.80 peak are observed for RNP1-3 cases. The broad distribution at lower FRET values suggests a heterogenous population and possibly some underlying dynamics. The smFRET traces shown in Fig. 2E-F show examples of static and dynamic cases, respectively. Additional traces are provided in Fig. S2. The fraction of dynamic traces was higher in RNP3 (∼30%) compared to RNP1 (∼5%) and RNP2 (∼10%) cases. The RNP3 case, which covers the entire VEGF-rQS, shows the greatest low FRET population, which is consistent with more efficient destabilization of the GQ and also the more frequent dynamic traces. On the other hand, for RNP4, the population at EFRET=0.80 remains unchanged, suggesting the GQ remains intact and the donor-acceptor separation does not change significantly. This could be due to inability of RNP4 to bind to its target in this truncated construct. The low FRET populations in RNP1-3 cases are in general modest in all tested conditions compared to the effects observed in other assays (reported in Fig. 3-6). This might be due to the smFRET experiments being performed at 25 °C, unlike all other measurements which were performed at 37 °C. Nevertheless, these studies illustrate structural changes being induced in this truncated VEGF mRNA construct when the VEGF-rQS is targeted.

**Figure 3.**
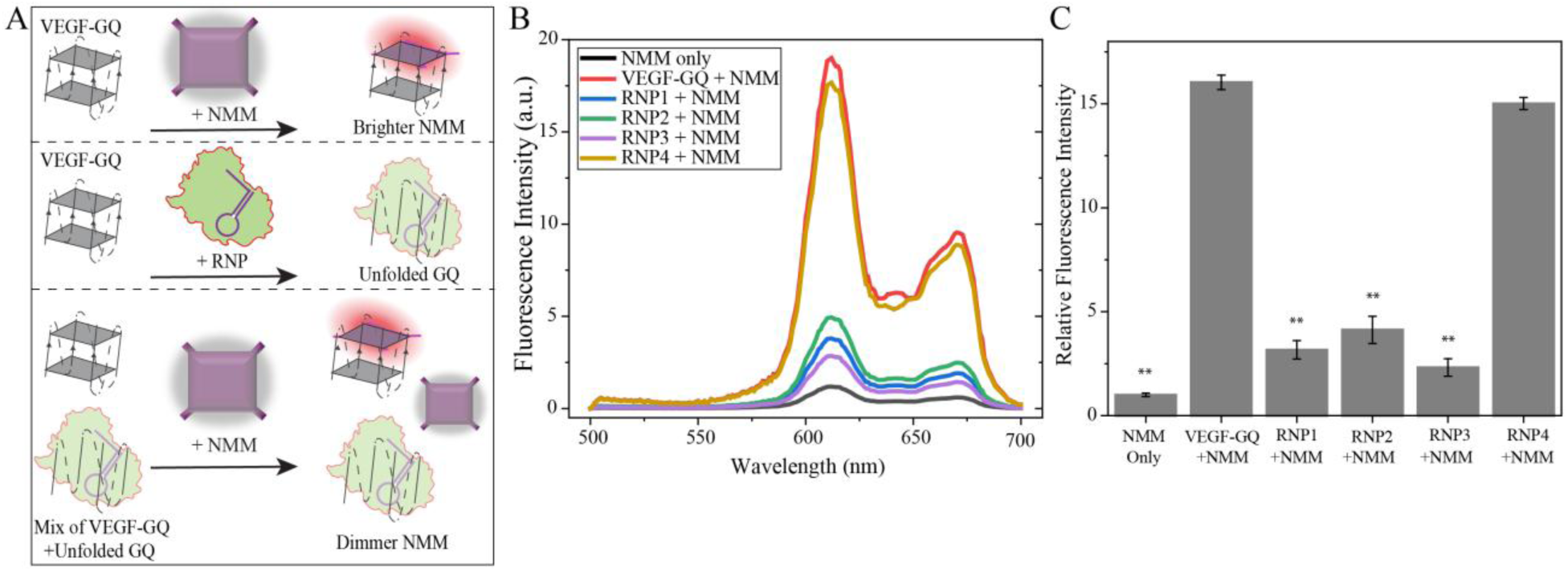
NMM fluorescence enhancement assay to probe impact of CRISPR-dCas13d binding on GQ stability. A) Schematic illustrating enhancement in NMM fluorescence upon binding to GQ. Destabilization of the GQ by CRISPR-dCas13d binding reduces the observed fluorescence enhancement. B) Fluorescence emission spectra of NMM before and after introducing RNA or RNP complexes. Binding to GQ enhances the emission intensity while different RNP complexes diminish this enhancement at different levels. RNP4, which does not bind to the VEGF-rQS, does not impact the fluorescence enhancement. C) Relative fluorescence emission intensity at 610 nm (average of at least 3 measurements) as normalized with respect to the NMM-only case. The error bars are the standard errors associated with the replicates. The ** symbols in (C) indicate a statistically significant difference compared to the VEGF-GQ + NMM case with p < .001. Table S2 lists t-test results of each comparison group.

To investigate GQ unfolding through an orthogonal approach, we employed the fluorescence enhancement of N-Methylmesoporphyrin IX (NMM) upon binding to GQ as a reporter. The premise of these bulk fluorescence measurements is that destabilization of the GQ by CRISPR-dCas13d RNP complexes should decrease NMM binding to GQ and hence reduce the fluorescence enhancement (Fig. 3A). NMM fluorescence emission is characterized by two peaks at ∼610 nm and ∼670 nm. Fig. 3B shows fluorescence emission spectra of NMM before it is mixed with a VEGF-GQ construct, which shows low levels of fluorescence emission. Introducing an RNA construct that contains a folded GQ (150 mM KCl) results in ∼16-fold increase in fluorescence emission compared to NMM-only case (16.0 ± 0.3 vs. 1.0 ± 0.1 a.u.), as quantified by the amplitude of the peak at 610 nm. All error bars reported on these experiments are based on standard error of the mean of at least three replicates. This is consistent with binding of NMM to the parallel RNA GQ structure. Mixing the RNA construct with RNP1, RNP2, or RNP3 before they are mixed with NMM shows significantly lower fluorescence enhancement: 3.2-fold, 4.1-fold, and 2.3-fold respectively. These correspond to (5.1 ± 0.4)-fold, (3.9 ± 0.7)-fold, and (6.9 ± 0.4)-fold reduction, respectively, compared to the absence of RNP complexes (RNA+NMM). In case of RNP3 targeting (the most effective destabilization), only 14% of the initial fluorescence remained. On the other hand, upon targeting with RNP4, which does not necessarily prevent GQ formation, 93.6% of the initial fluorescence intensity was retained (15.0 ± 0.3 vs. 16.0 ± 0.3 a.u.). This is consistent with RNP4 not stably binding to its target site. These results are summarized in Fig. 3C and they further support specific destabilization of the VEGF GQ by RNP1-3 while RNP4 does not impact GQ stability or accessibility to NMM.

To test validity of our *in vitro* findings in cellular setting, we performed dual-luciferase reporter assay in HeLa cells (Fig. 4A-B), western blot in HUVECs (Fig. 4C-D), and immunofluorescence assay in HUVECs (Fig. 5). A dual-luciferase reporter construct was prepared in which the entire IRES-A sequence was placed just upstream of the firefly luciferase, while the Renilla luciferase was under the control of a CMV promoter (hVEGFBicis, Fig. 4A). Firefly luciferase expression mediated by the VEGF IRES-A was normalized to Renilla luciferase expression transcribed from a CMV promoter and translated in a cap-dependent manner (Fig. 4B) ^32^. HeLa cells were transfected with this bicistronic plasmid and treated with different RNP complexes. The baseline activity of the untreated control (transfection reagent only, TR Only) represents uninhibited translation (1.0). RNP1, RNP2, and RNP3 treatments strongly repress translation initiation, as evidenced by reduction of normalized activity to 0.43 ± 0.06, 0.39 ± 0.06, and 0.34 ± 0.05, respectively. In contrast, RNP4 had little effect and kept luciferase activity at 97.3% ± 4.3% of the control, suggesting that it does not affect ribosome recruitment to IRES-A in cellular setting. These results demonstrate that RNP1, RNP2, and RNP3 destabilize the GQ structure and block ribosome recruitment to IRES-A.

**Figure 4.**
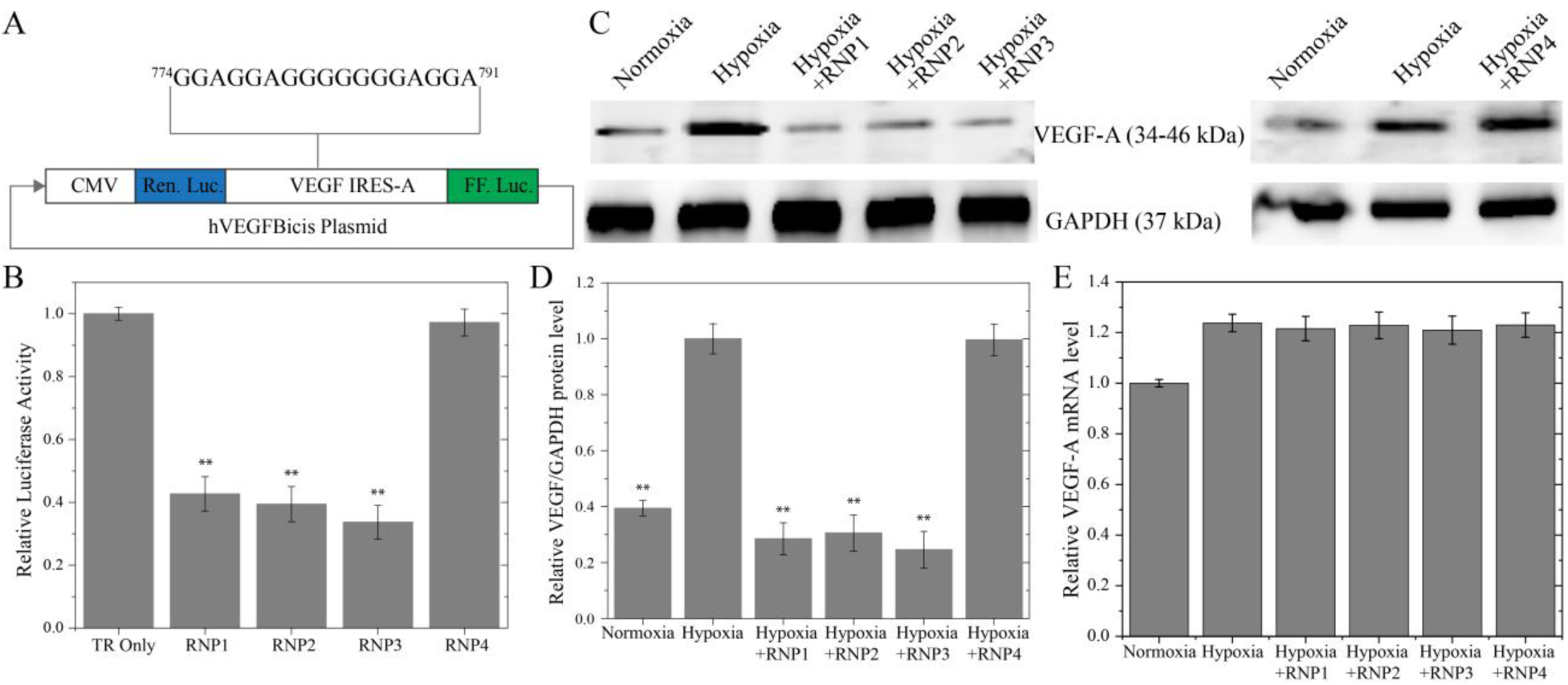
(A) Schematic of dual-luciferase bicistronic construct. Plasmids were transfected into untreated HeLa cells or those treated with different CRISPR-dCas13d complexes. (B) Histogram showing relative activity of the RNP1-4 complexes with respect to Transfection Reagent (TR) treated cells. (C) Images of western blot, where HUVECs were incubated with SNP to induce hypoxia and simultaneously treated with different CRISPR-dCas13d RNP complexes. D) Quantification of relative VEGF-A protein levels, where GAPDH was the internal control. (E) RT-qPCR studies illustrating VEGF mRNA levels before and after inducing hypoxia and treating with RNP1-4. The VEGF mRNA levels do show a significant change for RNP1-4 under hypoxia. Data bars represent means of independent experiments (n = 3), and the error bars are the standard errors. The ** symbols in (B) & (D) indicate a statistically significant difference compared to the Hypoxia case with p < .001. Table S4 lists t-test results of each comparison group for Fig. 4B, Table S5 for Fig. 4D, and Table S6 for Fig. 4E.

**Figure 5.**
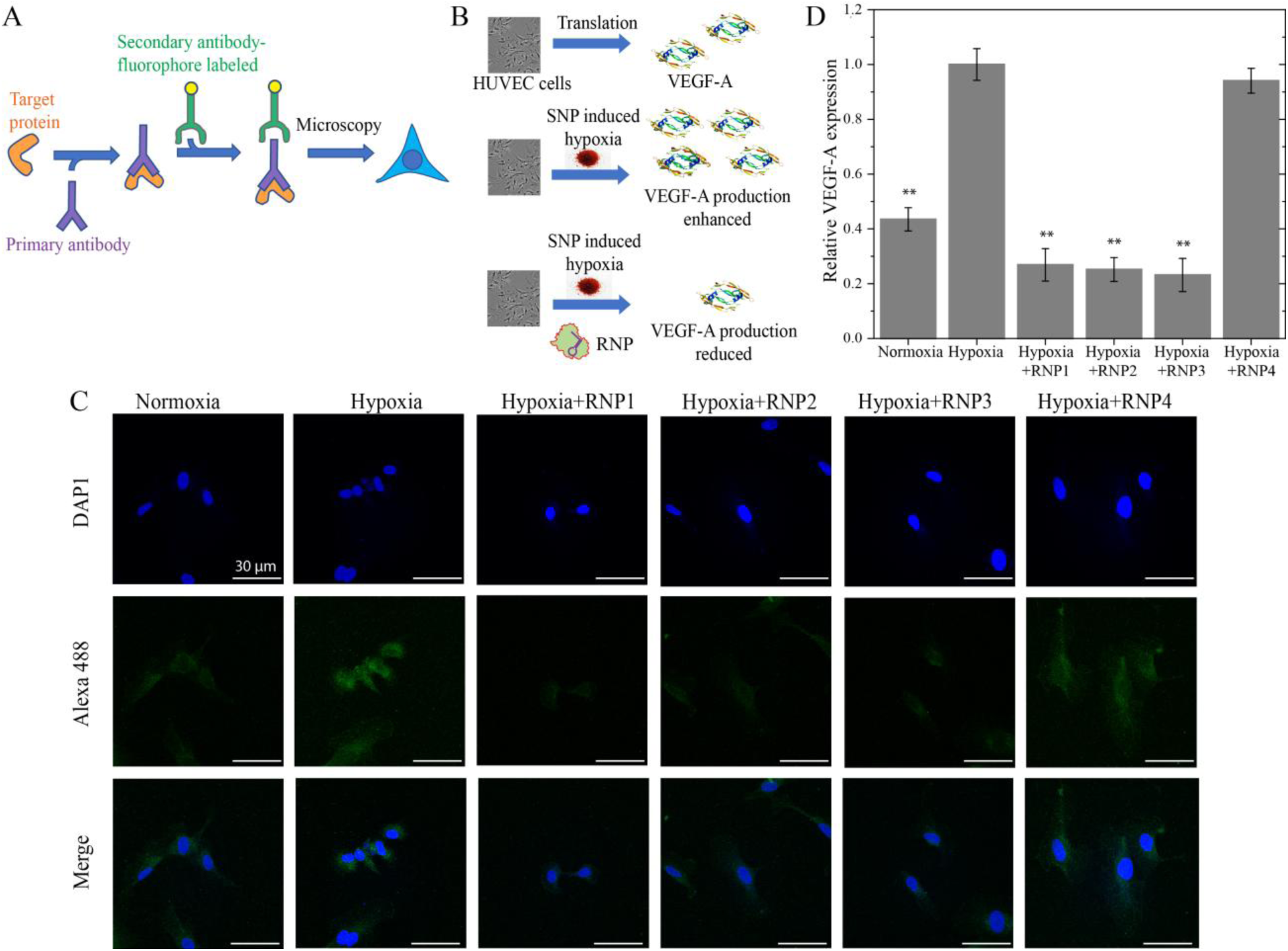
Immunofluorescence analysis of VEGF-A protein expression. A) Overview of the immunofluorescence assay. B) A pictorial summary of the concepts driving these measurements. C) Representative confocal microscopy images show different expression levels of VEGF-A protein for different treatment conditions. After fixation, cells were stained with DAPI (Blue) nuclear staining reagent and secondary VEGF-A antibody was labeled with Alexa 488 (Green). Scale bar is 30 μm in all images. D) More than a hundred cells were analyzed and quantified, and the histogram shows the average VEGF-A expression in these cells. Error bars are standard errors. The ** symbols in (D) indicate a statistically significant difference compared to the Hypoxia case with p < .001. Table S7 lists t-test results of each comparison group.

**Figure 6.**
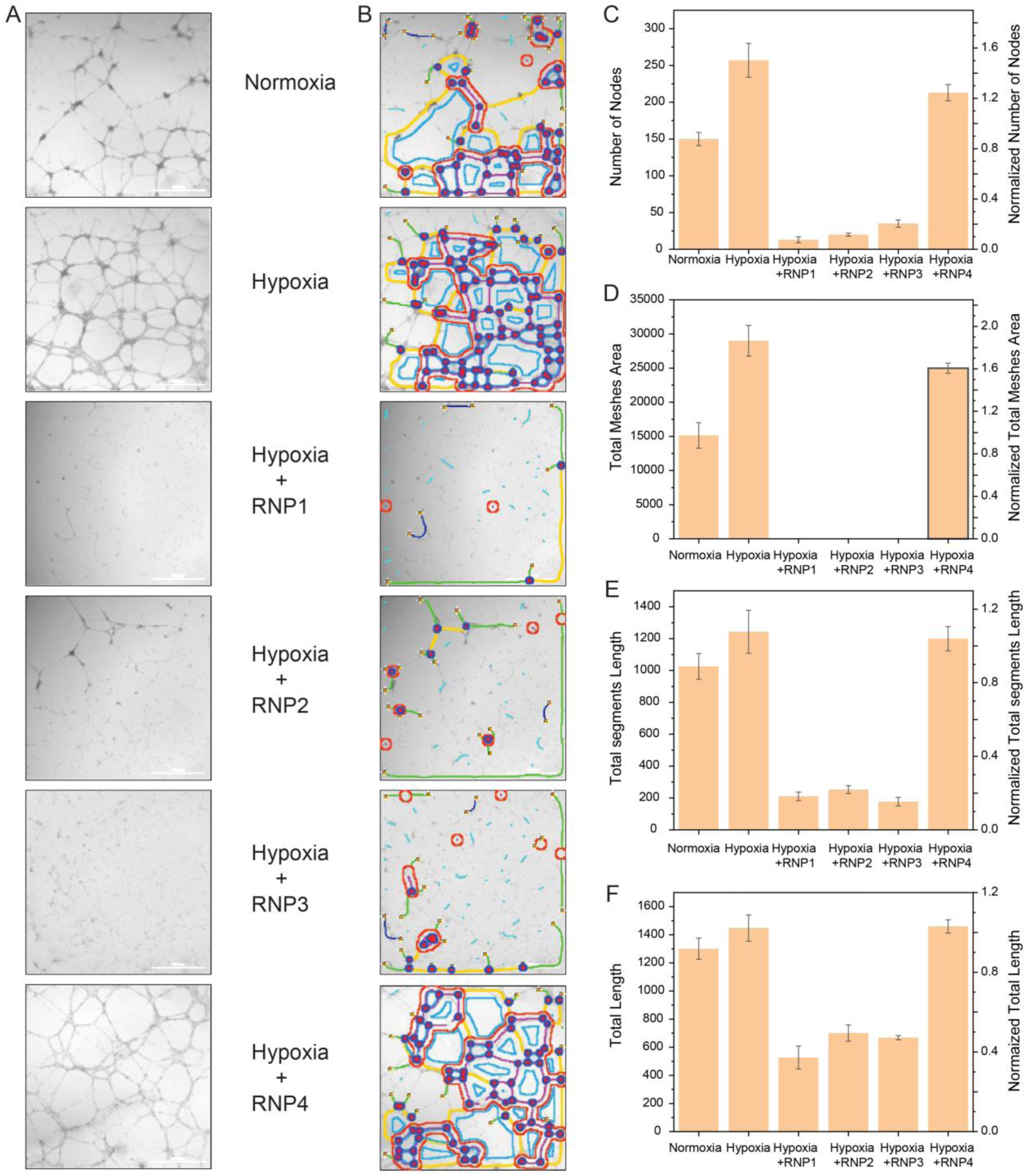
Quantification of antiangiogenic effect of CRISPR-dCas13d targeting using tube formation assay. Representative images of tube formation assay using Bovine Aortic Endothelial Cells (BAOECs) after 16 hours of treatment with different RNP complexes. (B) Analysis of tube formation assay using ImageJ software. Histograms show the quantification results for the following parameters. (C) Number of Nodes, (D) Total mesh area, (E) Total segment length, and (F) Total length (sum of Branches and segment length). Data are expressed as mean ± standard deviation of three independent experiments. Scale bar = 1000 µm. Table S7-S10 list t-test results of each comparison group for Fig. 6C-F, respectively.

For western blot analysis, we used human umbilical vein endothelial cells (HUVECs) and induced hypoxia using Sodium nitroprusside (SNP), which is a strong hypoxia mimic ^61^. The cells were treated with SNP and the same four CRISPR-dCas13d RNP complexes (RNP1–RNP4). Protein levels were normalized with respect to GAPDH (Fig. 4C). A baseline expression of VEGF-A (34-46 kDa band) was observed in normoxia (not treated with SNP). This expression increased by (2.5 ± 0.2)-fold upon treating cells with SNP, which was taken as a reference level to quantify the impact of RNP complexes. Introducing the RNPs under hypoxia (SNP+RNP) reduced VEGF-A expression by 3.5-fold, 3.3-fold and 4.1-fold for RNP1, RNP2, and RNP3, respectively (0.28 ± 0.05, 0.30 ± 0.06, 0.24 ± 0.06 where SNP-only case is normalized to 1.00). On the other hand, RNP4 was identical to the SNP-only case (0.99 ± 0.05). These results are summarized in Fig. 4D. Real time quantitative polymerase chain reaction (RT-qPCR) measurements show that VEGF mRNA levels are unaffected by RNP1-4 treatment (Fig. 4E) and that their effect is specific to translation level regulation.

Finally, we performed immunofluorescence measurements to quantify the impact of CRISPR–dCas13d RNPs on VEGF expression in HUVECs, which overexpress VEGF-A protein during hypoxic conditions. A 24-h SNP treatment was used to induce hypoxia. Cells that were treated with just SNP or SNP and an RNP complex were imaged using confocal microscopy. VEGF-A protein levels were quantified using an immunofluorescence assay (Fig. 5A-B). VEGF-A expression increased by 2.8 times in response to SNP treatment, indicating that hypoxia-driven upregulation of VEGF was successfully induced. Under hypoxic conditions, this increase was considerably reduced by treatment with GQ-targeting RNPs: RNP1 reduced VEGF-A expression by 3.7 ± 0.7-fold, RNP2 by 3.9 ± 0.6-fold, and RNP3 by 4.3 ± 0.9-fold compared to cells treated with SNP-only. RNP4 showed no discernible impact (only a ∼7% drop from hypoxia levels). Fig. 5C shows representative images and Fig. 5D shows a quantitation of the fluorescent marker levels.

Tube formation assay (Fig. 6 and Table 1) showed that targeting the VEGF-rQS using CRISPR-dCas13d under hypoxia significantly impacted endothelial tube development. These studies showed that angiogenesis significantly increased under hypoxia compared to normoxia as quantified in terms of node number (257 ± 23 vs. 150 ± 9), total mesh area (29007 ± 2250 vs. 1869 ± 1869), segment length (1243 ± 135 vs. 1026 ± 81), and total tube length (1448 ± 93 vs. 1301 ± 75). On the other hand, tube formation was significantly suppressed when the VEGF-rQS was targeted with dCas13d complexes (RNP1, RNP2, and RNP3) under hypoxia, which resulted in nearly total loss of mesh structures, an order of magnitude decrease in node numbers, and significant decreases in total segment length and total length. On the other hand, RNP4 complex, which does not target the VEGF-rQS, did not result in a significant change in angiogenic capacity, indicating that this guide does not inhibit VEGF-mediated angiogenesis.

**Table 1.** Mean and standard deviation of relevant parameters of tube formation assay (Fig. 6) for different CRISPR-dCas13d complexes.

| Rd13 Complex | Nb of nodes | Tot. meshes area | Tot. segments length | Tot. length |
| --- | --- | --- | --- | --- |
| Normoxia | 150±9 | 15141±1869 | 1026±81 | 1301±75 |
| Hypoxia | 257±23 | 29007±2250 | 1243±135 | 1448±93 |
| Hypoxia+RNP1 | 13±4 | 0±0 | 211±27 | 527±81 |
| Hypoxia+RNP2 | 20±2 | 0±0 | 253±25 | 701±58 |
| Hypoxia+RNP3 | 35±5 | 0±0 | 177±27 | 668±15 |
| Hypoxia+RNP4 | 213±11 | 24983±738 | 1200±76 | 1460±47 |

## Conclusion

This research uses the CRISPR-dCas13d system to target functional RNA structures, establishing a novel mechanism for translation level gene expression regulation. Using an array of *in vitro* biochemical, cellular, and functional assays, we have shown that CRISPR-dCas13d can be used to destabilize a translationally-active G-quadruplex structure, which leads to repression of VEGF expression. By using nuclease-dead dCas13d, our approach avoids irreversible genomic editing, which also ameliorates off-target effects, and provides a transient and reversible therapeutic intervention. This approach offers a major benefit over small molecule inhibitors, which can interfere with a variety of biological processes as they frequently lack selectivity. Our results show the feasibility of blocking cap-independent translation, which is a mechanism often used by cancer cells to maintain survival and growth under stress conditions. Finally, our approach provides an adaptable platform to target other genes with translationally active secondary structures, potentially serving as an effective RNA-based therapeutic.

### Supporting Information Available

Sequences of DNA constructs, tables listing results of statistical analysis demonstrating significance of observed variations.

## AUTHOR INFORMATION

### Corresponding Author

* Hamza Balci − Department of Physics, Indiana University, Indianapolis, IN 46202, USA;

* Soumitra Basu − Department of Chemistry and Biochemistry, Texas State University, San Marcos, TX 78666, United States;

## Author CRediT Statement

**Mohammad Lutful Kabir:** Conceptualization, Investigation, Formal Analysis, Writing-Original Draft. **Sineth G. Kodikara:** Investigation, Visualization. **Najmah Al Ramel:** Investigation. **Janan Alfehaid:** Investigation, Visualization. **Soumitra Basu:** Supervision, Funding Acquisition, Writing-Review & Editing, **Hamza Balci:** Supervision, Funding Acquisition, Writing-Original Draft

## Conflict of Interest Statement

The authors declare that they have no known competing financial interests or personal relationships that could have appeared to influence the work reported in this paper.

### Data Availability Statement

The data supporting the findings of this study are available from the corresponding author upon reasonable request.

## Acknowledgements

This study was supported by the National Institutes of Health under Grant No. 1R35GM156183 to H.B. and 1R15GM146180 to H.B. and S.B. J.A. is funded by the Deanship of Scientific Research at Northern Border University, Arar, KSA through the project number NBU-SAFIR-2026.

