## Supplementary Information for "Cap-Independent Translation Regulation of VEGF Using CRISPR-dCas13d"

### Sequences of Nucleic Acids

The sequences of the nucleic acids used in the manuscript are listed in Table S1, along with the particular figures that were used. The VEGF-G-quadruplex sequence (VEGF-rQS) is marked with red fonts in the mRNA strand. The 5′ Cy3-labeled nucleotide is shown in green in VEGF-mRNA. The 22 nt-long guide sequences are marked with purple fonts in all the guide RNA (gRNA) sequences. The Bottom 12 DNA strand is complementary to a segment at the 3'-end of VEGF-mRNA and is labeled with Cy5.

**Table S1:** *Oligonucleotide sequences used for in vitro and in vivo assays.*

| Name | Sequence (5' to 3') | Figure |
|---|---|---|
| **VEGF-mRNA labeled** | UGGAGGAGCCGCAGCCGGAGGAGGGGGAGGAGGAAGAAGAGAAGGA AUGGCGACGGCAGC | Fig. 2 |
| **VEGF-mRNA non-labeled** | UGGAGGAGCCGCAGCCGGAGGAGGGGGAGGAGGAAGAAGAGAAGGA AUGGCGACGGCAGC | Fig. 3 |
| **gRNA-1** | CACUAGUGCGAAUUUGCACUAGUCUAAAACCCCCCUCCUCCGGCUGC GGCUC | Fig. 2-6 |
| **gRNA-2** | CACUAGUGCGAAUUUGCACUAGUCUAAAACCCUUCUCUUCUUCCUCC UCCCC | Fig. 2-6 |
| **gRNA-3** | CACUAGUGCGAAUUUGCACUAGUCUAAAACUCUUCCUCCUCCCCCUCC UCCG | Fig. 2-6 |
| **gRNA-4** | CACUAGUGCGAAUUUGCACUAGUCUAAAACUCCGGCUGCGGCUCCUC CAGG | Fig. 2-6 |
| **Bottom 12** | /5Biosg/ GCT GCC GTC GCC / 3Cy5sP/ | Fig. 2 |
| **GAPDH-Forward Primer** | AGGTCGGTGTGAACGGATTTG | Fig. 4 |
| **GAPDH-Reverse Primer** | GGGGTCGTTGATGGCAACA | Fig. 4 |
| **VEGF Forward Primer** | TTGCCTTGCTGCTCTACCTCCA | Fig. 4 |
| **VEGF Reverse Primer** | GATGGCAGTAGCTGCGCTGATA | Fig. 4 |

Table S2: T-test analysis on the data presented in Figure 3C.

| Name | T- Test | Comment |
|---|---|---|
| NMM Only | t (2) = -130.18, p < 0.00001 | Significantly different |
| RNP1+NMM | t (2) = -18.64, p < 0.0001 | Significantly different |
| RNP2+NMM | t (2) = -17.39, p < 0.0001 | Significantly different |
| RNP3+NMM | t (2) = -19.91, p < 0.0001 | Significantly different |
| RNP4+NMM | t (2) = -2.94, p > 0.05 | No Significant difference |

Table S3: T-test analysis on the data presented in Figure 4B.

| Name | T- Test | Comment |
|---|---|---|
| RNP1 | t (2) = -18.15, p < 0.0001 | Significantly different |
| RNP2 | t (2) = -18.65, p < 0.0001 | Significantly different |
| RNP3 | t (2) = -20.37, p < 0.0001 | Significantly different |
| RNP4 | t (2) = -0.93, p < 0.0001 | No Significant difference |

Table S4: T-test analysis on the data presented in Figure 4D.

| Name | T- Test | Comment |
|---|---|---|
| Normoxia | t (2) = 17.60, p < 0.0001 | Significantly different |
| Hypoxia+RNP1 | t (2) = 15.90, p < 0.0001 | Significantly different |
| Hypoxia+RNP2 | t (2) = 14.51, p < 0.0001 | Significantly different |
| Hypoxia+RNP3 | t (2) = 15.63, p < 0.0001 | Significantly different |
| Hypoxia+RNP4 | t (2) = 0.91, p = 0.719 | No Significant difference |

Table S5: T-test analysis on the data presented in Figure 4E.

| Name | T- Test | Comment |
|---|---|---|
| SNP | t (2) = 10.21, p < 0.0001 | Significantly different |
| RNP1 | t (2) = -0.76, p = 0.477 | No Significant difference |
| RNP2 | t (2) = 0.27, p = 0.798 | No Significant difference |
| RNP3 | t (2) = -0.70, p = 0.509 | No Significant difference |
| RNP4 | t (2) = -0.27, p = 0.798 | No Significant difference |

Table S6: T-test analysis on the data presented in Figure 5D.

| **Name** | **T- Test** | **Comment** |
|---|---|---|
| Normoxia | $t(2) = -14.60, p < 0.0001$ | Significantly different |
| Hypoxia+RNP1 | $t(2) = -15.58, p < 0.0001$ | Significantly different |
| Hypoxia+RNP2 | $t(2) = -20.86, p < 0.0001$ | Significantly different |
| Hypoxia+RNP3 | $t(2) = -14.90, p < 0.0001$ | Significantly different |
| Hypoxia+RNP4 | $t(2) = -1.24, p = 0.272$ | No Significant difference |

Table S7: T-test analysis on the data presented in Figure 6C.

| **Name** | **T- Test** | **Comment** |
|---|---|---|
| Normoxia | $t(2) = 3.44, p = 0.03$ | Significantly different |
| Hypoxia+RNP1 | $t(2) = 7.44, p = 0.008$ | Significantly different |
| Hypoxia+RNP2 | $t(2) = 7.25, p = 0.009$ | Significantly different |
| Hypoxia+RNP3 | $t(2) = 6.80, p = 0.01$ | Significantly different |
| Hypoxia+RNP4 | $t(2) = 1.63, p = 0.12$ | No Significant difference |

Table S8: T-test analysis on the data presented in Figure 6D.

| **Name** | **T- Test** | **Comment** |
|---|---|---|
| Normoxia | $t(2) = 6.70, p = 0.001$ | Significantly different |
| Hypoxia+RNP1 | $t(2) = 18.22, p = 0.001$ | Significantly different |
| Hypoxia+RNP2 | $t(2) = 18.22, p = 0.001$ | Significantly different |
| Hypoxia+RNP3 | $t(2) = 18.22, p = 0.001$ | Significantly different |
| Hypoxia+RNP4 | $t(2) = 2.40, p = 0.06$ | No Significant difference |

Table S9: T-test analysis on the data presented in Figure 6E.

| **Name** | **T- Test** | **Comment** |
|---|---|---|
| Normoxia | $t(2) = 1.94, p = 0.07$ | No Significant difference |
| Hypoxia+RNP1 | $t(2) = 10.54, p = 0.004$ | Significantly different |
| Hypoxia+RNP2 | $t(2) = 10.15, p = 0.004$ | Significantly different |
| Hypoxia+RNP3 | $t(2) = 10.90, p = 0.004$ | Significantly different |
| Hypoxia+RNP4 | $t(2) = 0.39, p = 0.36$ | No Significant difference |

Table S10: T-test analysis on the data presented in Figure 6F.

| **Name** | **T- Test** | **Comment** |
|---|---|---|
| Normoxia | t (2) = 1.73, p = 0.07 | No Significant difference |
| Hypoxia+RNP1 | t (2) = 10.47, p = 0.0002 | Significantly different |
| Hypoxia+RNP2 | t (2) = 9.57, p = 0.001 | Significantly different |
| Hypoxia+RNP3 | t (2) = 11.66, p = 0.003 | Significantly different |
| Hypoxia+RNP4 | t (2) = 0.17, p = 0.43 | No Significant difference |

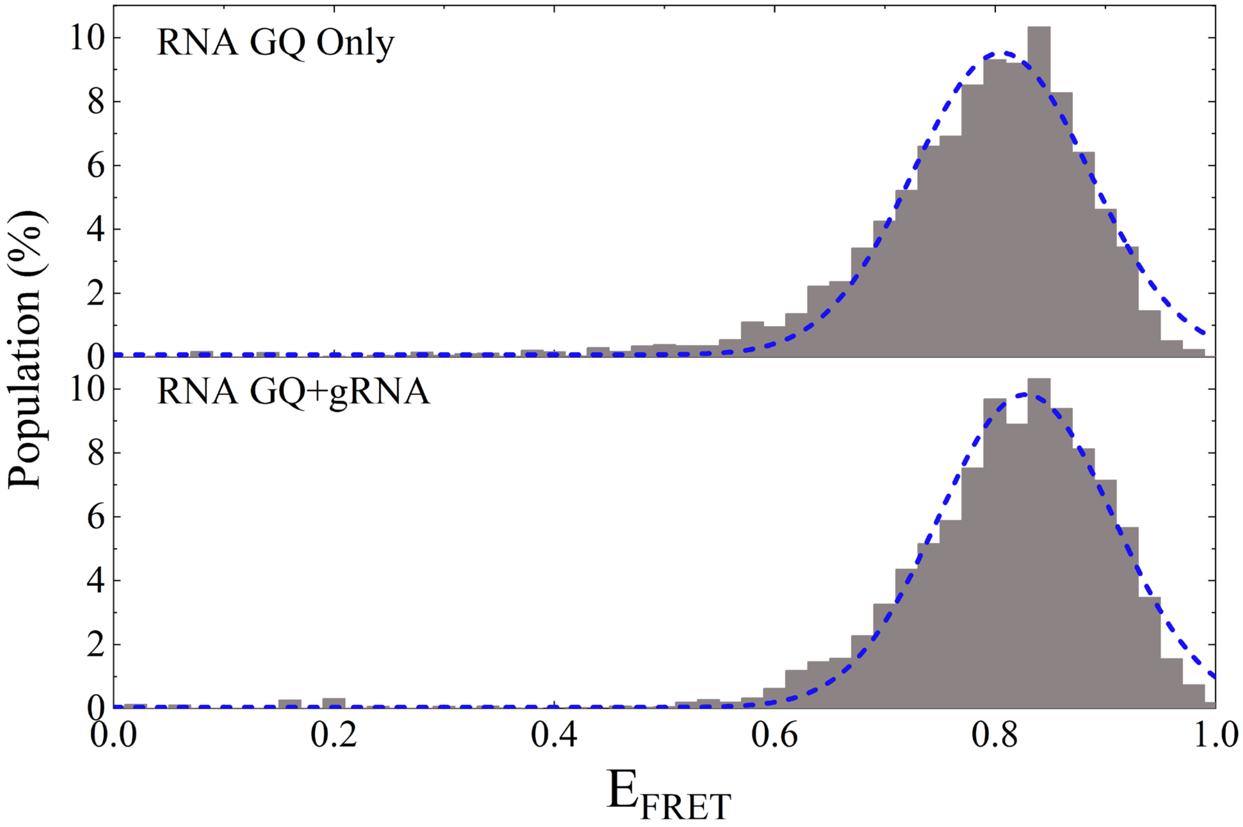


Figure S1: Single molecule FRET histograms demonstrating the inability of guide RNA to unfold the VEGF GQ in the absence of dCas13d.

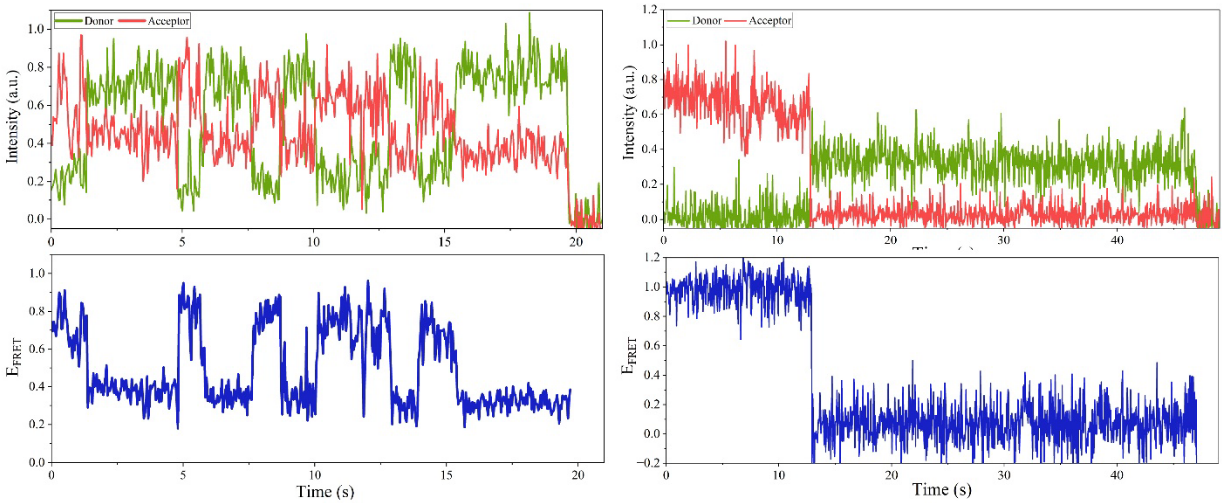


Figure S2: Additional smFRET traces showing dynamic (left) and static (right) FRET levels.
